# The illusion of uniformity does not depend on low-level vision: evidence from sensory adaptation

**DOI:** 10.1101/278721

**Authors:** Marta Suárez-Pinilla, Anil K. Seth, Warrick Roseboom

## Abstract

Visual experience appears richly detailed despite the poor resolution of the majority of the visual field, thanks to foveal-peripheral integration. The recently described Uniformity Illusion (UI), in which peripheral elements of a pattern seem to take on the properties of foveal elements, may shed light on this integration. We examined the basis of UI by generating adaptation to a pattern of Gabors suitable for producing UI on orientation. After removing the pattern, participants reported the tilt of a single peripheral Gabor. The tilt after-effect (TAE) followed the physical adapting orientation rather than the global orientation perceived under UI, even when the illusion had been reported for a long time. Conversely, a control experiment replacing illusory for physical uniformity for the same durations did produce an after-effect to the global orientation. Our results indicate that the UI is not associated with changes in sensory encoding, but likely depends on high-level processes.

## INTRODUCTION

Visual experience appears richly detailed despite the poor sensory precision of the majority (periphery) of the visual field. This topic has received considerable recent attention [1, 2], with debate about the degree to which visual experience is in fact rich, and the potential perceptual processes that may contribute to apparent richness. One recent study demonstrated a compelling example of how the rich detail within the high-precision central visual field alters peripheral perception - the Uniformity Illusion (UI) [3]. UI describes a phenomenon wherein apparent perceptual uniformity occurs when variable sensory stimulation is presented in peripheral vision, while the central visual field is presented with uniform stimuli. UI occurs for a wide variety of perceptual dimensions, including relatively low-level sensory features like orientation or colour, and higher-level features such as density (see www.uniformityillusion.com for examples).

We sought to examine the mechanisms underlying UI using perceptual adaptation. It is well established that exposure to a specific stimulus (like an oriented grating) causes perceptual after-effects (e.g. tilt after-effect; **TAE**) [4]. For visual orientation, perceptual after-effects have been associated with specific changes in neural coding and are localised in a retinotopic reference frame [5]. Here, we utilise the spatial specificity of TAE to examine whether the apparent perceptual uniformity in UI can be attributed to changes in low-level neural coding for visual orientation. Specifically, we presented participants with Gabor grids wherein the orientation of central elements was uniform, but the orientation of peripheral elements was variable -producing UI. At fixed test locations in the periphery of the grid, we presented a physical orientation that differed from the global illusory percept, thus putting local and global orientation in opposition. Following prolonged exposure to global illusory uniformity (UI), we contrasted whether the resultant TAE was consistent with the local, physical orientation or with the illusory global orientation.

## METHODS

### Procedure

The experiment had two parts: Illusion session and Control session. Each session contained six blocks, and each block had an adaptation phase and a test phase (Figure 1). A practice block was run before the Illusion session to familiarise participants with UI.

**Figure 1.**
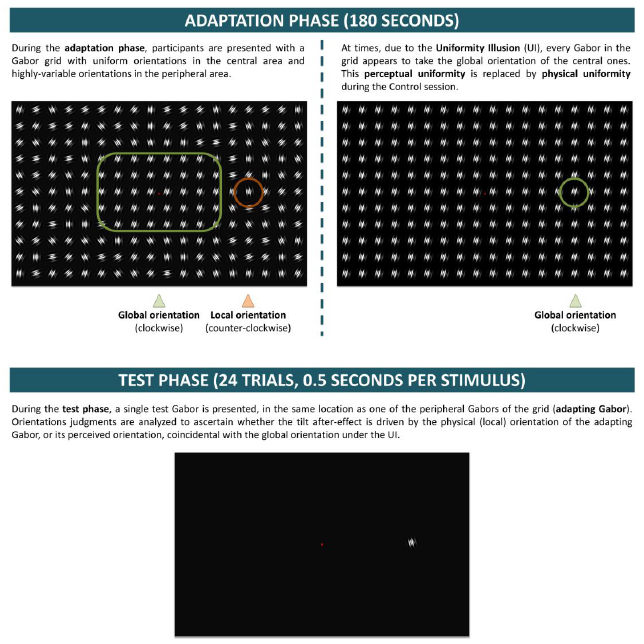
Experimental structure. During the adaptation phase, participants were presented with a Gabor grid wherein the central Gabors had a uniform orientation, while peripheral orientations were heterogeneous. Under UI, perceptual experience was that of a uniform pattern with all Gabors tilted like the central ones. This illusory percept alternated with a non-illusory, non-uniform percept at different times during adaptation. For a specific peripheral Gabor (adapting Gabor), physical and illusory orientation were always in opposition. The Control session replicated the phenomenology of the Illusion session, replacing perceived with physical uniformity at times in which the participant reported UI in the Illusion session. The test phase had 24 trials, wherein participants reported the tilt of a single peripheral Gabor whose location coincided with the adapting Gabor.

### Illusion session

Each block began with an adaptation phase, in which participants were presented with a grid of Gabor patches suitable for producing the UI, affecting the apparent orientation of peripheral elements: all Gabors in the central area had a uniform orientation, whereas orientation of the peripheral Gabors was heterogeneous. Gaze-contingent stimulus presentation ensured that each Gabor was presented to a specific retinal location, as the entire pattern was removed if the participant’s gaze deviated from central fixation by more than 1.5 degrees of visual angle (dva).

Adaptation lasted 180 seconds but, because the stimulus was removed when fixation lapsed, actual exposure time could be shorter.

Participants reported the experience of illusory uniformity by pressing a key when all Gabors appeared to take a uniform orientation.

The test phase had 24 trials, separated by a pseudo-random interval of 1000-1500 ms. In each trial, a single Gabor (test Gabor) was presented for 500 ms at a specific peripheral location, coinciding with the position of a specific Gabor during adaptation (adapting Gabor). Participants reported if the test Gabor was tilted clockwise (CW) or counter-clockwise (XCW) from vertical.

### Control session

The Control session also had six blocks, each built to replicate the phenomenology of a homologous block of the Illusion session but replacing illusory for physical uniformity during the adaptation phase.

During the adaptation phase in the Illusion session, an empty background was presented whenever the gaze-contingent mechanism removed the adapting pattern. The same pattern of stimulus presentation and removal was replicated in the Control session. The stimulus was additionally removed whenever fixation lapsed in the Control session. At any other time, the presentation displayed one of two patterns, differing only in the orientation of peripheral Gabors. The first was identical to the pattern presented in the Illusion session and was displayed at times in which the participant had *not* reported UI during adaptation in the Illusion session. At times during which the participant had reported UI, the presented pattern was one in which all Gabors had the same *physical* orientation, consistent with the desired illusory orientation during the Illusion session. Thus, physical uniformity was inserted at the times in which illusory uniformity had been reported in the Illusion session. Participants were not informed that this would occur.

The test phase was identical to that in the Illusion session: the location and orientation of the test Gabor in each trial was identical, as well as its test latency (time between the end of the adaptation phase and stimulus onset).

### Stimuli

Stimuli were displayed on dark grey background (1.96 cd/m^2^). A red fixation dot (8.34 cd/m^2^, 0.42 dva diameter) showed constantly on the screen centre.

### Gabor patches

Each Gabor consisted of a sine-wave luminance grating with Michelson contrast of 1, 0° phase and spatial frequency of 1.66 cycles per dva (cpd), and a 2-D Gaussian envelope with a sigma of 0.43 dva.

### Adapting pattern

The adapting pattern spanned the entire screen and consisted of a 13×17 grid formed by invisible square cells measuring 3 dva per side (Figure 1). Each Gabor was presented in the centre of each cell. The central area spanned 15 dva horizontally and vertically, encompassing all cells belonging to rows 5-9 and columns 7-11. Accordingly, Gabors were classified into central and peripheral, which determined their orientation. All central Gabors had the same orientation, which could be one of two values, each for half the blocks of one session: −15°(global clockwise tilt, GCW) or 15°(global counter-clockwise tilt, GXCW). The orientations of peripheral Gabors were sampled from a discrete uniform distribution centred on the global orientation and ranging 70°(35°to each side). Thus, mean orientation was the same for central and peripheral Gabors and matched the global orientation perceived under UI.

Two peripheral Gabors of the pattern (adapting Gabors) corresponded to the positions in which the test Gabors would be displayed during the test phase: they were located along the middle (7^th^) row, at 12.02 dva left and right of the screen centre (columns 5 and 13). Both had the same non-randomized local orientation, which was the opposite of the global orientation of the block: either 15°(local counter-clockwise tilt, LXCW) or −15°(local clock-wise, LCW).

Henceforth we give the label **adapting condition CX** to the presentation pattern wherein the local orientation of the adapting Gabor is clockwise and the global orientation of the pattern is counter-clockwise (LCW, GXCW). Conversely, we will refer to the pattern with LXCW and GCW orientations as **adapting condition XC**. Both conditions occurred equally frequently during the experiment.

As described above, during the Control session, the adapting pattern was replaced by a physically uniform pattern at those times during which participants had reported UI in the Illusion session. In these instances, *every* Gabor in the pattern (including the adapting Gabors) took the global orientation.

### Test Gabors

A single test Gabor was presented per trial, matching the position of one of the two adapting Gabors. Test Gabors were displayed in the left and right hemifield with equal frequency per block and could take one of eight equally frequent orientations: −12°, −5°, −2°, −1°, 1°, 2°, 5°and 12°(negative values indicate clockwise tilt). Thus, test orientations were always intermediate between global and local orientations (−15°, 15°).

### Participants

Participants were recruited through online advertisement, over 18 and reported normal or corrected-to-normal vision. This study received ethical approval by the Research Ethics Committee of the University of Sussex.

### Apparatus

Experiments were programmed in MATLAB 2016a (MathWorks Inc., Natick, US-MA) and displayed on a LaCie Electron 22BLUE II 22‘’ with screen resolution of 1024×768 pixels and refresh rate of 100 Hz. Eye-tracking was performed with Eyelink 1000 Plus (SR Research, Mississauga, Ontario, Canada) at sampling rate of 1000 Hz, with level desktop camera mount. Head position was stabilized 43 cm from the screen using chin and forehead rest.

### Statistical analysis

Statistical analyses were conducted using Matlab 2017b (with Palamedes toolbox, version 1.8.1 for Psychometric curve fitting) and JASP (JASP Team (2017), version 0.8.3.1). For Bayesian t-tests we employed as prior distribution Cauchy(0, 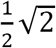) for two-sided predictions, or a folded Cauchy-(0, 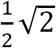) for one-sided predictions (measure 1 < measure 2). Likewise, for Bayesian Pearson correlations we employed a uniform distribution U(−1,1) or U(0,1) for two-sided or one sided (positive) predictions, respectively. For each analysis, the prior utilised is indicated by the formulated prediction and the subscripts in BF10 (two-sided) or BF-0 /BF+0 (one-sided).

## RESULTS AND DISCUSSION

Thirty participants volunteered for the experiment: 23 female, mean age 21.6.

To ensure sufficient exposure to the adapting pattern, we excluded blocks wherein the pattern had been displayed for less than 2/3 of the adaptation phase (<120 seconds), due to gaze-contingent stimulus removal. In such cases, the corresponding blocks from both Control and Illusion sessions were removed, to maintain balance. This caused exclusion of 32.78% blocks (118/360), including the entire datasets from five participants. Results presented here correspond to the blocks of the remaining twenty-five participants.

### Adaptation phase

Average exposure time to the adapting pattern per block was 164.43 and 149.57 seconds for the Illusion and Control sessions: 91.35% and 83.09% of the adaptation phase, respectively. The lower proportion in the Control session was expected as pattern removal occurred whenever it had in the Illusion session, in addition to times of improper fixation in the Control block.

Perceived uniformity was reported, on average, for 43.87 seconds in the Illusion session, comprising 26.92% of the time of pattern presentation (minimum 0.55%, maximum 72.23%). The proportion of time of perceived uniformity during the Control session was no different than for the Illusion session: 28.25% (minimum 0.59%, maximum 78.42%, Bayesian paired-samples t-test: BF01=3.207 -moderate evidence for the null hypothesis). Physical uniformity in the Control session was reported as perceptually uniform 68.11% of the time; by contrast, the non-uniform pattern was reported as uniform only 8.58% of the time. Possibly, presentation of a truly uniform pattern shifted a subjective criterion for uniformity, leading to more conservative reports in the Control sessions.

### Hypotheses and measurements

The experiment placed adaptation to illusory and physical orientation in opposition to disambiguate two competing hypotheses:

1. The perceived orientation under UI has no effect on tilt adaptation; the TAE is driven solely by the physical orientation of the adapting Gabor.
2. The global orientation perceived for the entire pattern (including the adapting Gabor) under UI can produce a TAE.

To decide between hypotheses, data was analysed to ascertain the direction of the adaptation-induced bias. We calculated the proportion of XCW reports per test Gabor orientation and obtained the best-fitting cumulative Gaussian psychometric curve. The point of subjective equality (PSE) was defined as the test orientation at which 50% reports are XCW. Since CW orientations have (conventionally) negative sign and vice versa, negative PSE indicates a XCW bias and positive PSE a CW bias.

During the Illusion session, a TAE driven by (i.e. away from) the local orientation of the adapting Gabor would imply physical adaptation, while a global-driven TAE would indicate adaptation to illusory orientation. During the Control session, both local and global-driven TAE are compatible with physical adaptation, since the adapting Gabor physically takes the global orientation at times of reported illusory uniformity in the Illusion session.

By calculating participants’ PSE per adapting condition we obtained two measurements:

1. **PSECX** and **PSEXC.** For a local-driven TAE, responses for adapting condition CX should exhibit a XCW bias compared to condition XC (PSECX<PSEXC), and the reverse should happen for a global-driven TAE.
2. **dPSE**=PSECX–PSEXC. We employ this as a summary measure indicating the overall direction of the bias. A **negative dPSE** indicates a predominance of local-driven TAE (PSECX<PSEXC) consistent with **physical adaptation** to the local orientation, while a **positive dPSE** indicates a global-driven TAE, consistent with **adaptation to the illusion** (or to the physical replication of the illusion during the Control session).

## The TAE is driven by physical orientation, not by illusory orientation

### Overall effect

#### Illusion session

Figure 2A presents the psychometric curves for a representative participant’s reports during the Illusion session, separated by adapting condition (CX or XC). Figure 2B presents the average PSE across 25 participants, per condition: PSECX=-0.477o, PSEXC=0.607°, indicating a XCW and CW bias, respectively. On average, dPSE=-1.201° (PSECX<PSEXC Bayesian paired-samples t-test: BF-0=3.078) indicated a local, physical-driven adaptation.

**Figure 2.**
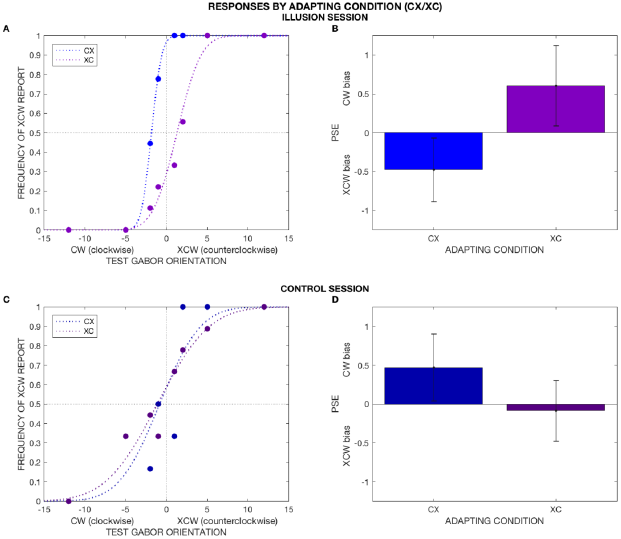
Response patterns by adapting condition. Figures **2A** and **2C** present a representative participant’s proportion of counter-clockwise (XCW) reports per test Gabor orientation, separated by adapting condition, during the Illusion (2A) and Control (2C) sessions. The dotted lines show the best cumulative Gaussian fit for the psychometric curve of each condition. Figures **2B** and **2D** depict the average point of subjective equality (PSE) for the entire sample (N=25), separated by condition; the error bars show the between-subject standard error. A negative PSE indicates a XCW bias and a positive PSE a CW bias. **2A-2B.** Illusion session. PSEs for both adapting conditions reflect a bias *away* from local orientation (local-driven TAE). **2C-2D.** Control session. On average (2D) responses show a global-driven TAE in CX and are unbiased in XC, although in the single participant’s example (2C), both conditions exhibit a XCW bias, likely a systematic individual bias. These results show that perceived (illusion) and physical (control) uniformity behave differently, suggesting that the TAE is always driven by the physical orientation, even when that orientation is unseen under UI.

#### Control session

In the Control session, the adapting Gabor physically takes the global orientation of the pattern during times of reported uniformity in the Illusion session. If adaptation is produced by physical orientation, we should observe a more global-driven TAE compared to the Illusion session: dPSEIL<dPSECO. Conversely, if perceived orientation under UI causes adaptation, we should not see a difference between perceived and physical uniformity: dPSEIL=dPSECO.

Results indicate predominance of global-driven TAE during the Control session (Figure 2D): PSECX-CO=0.607°, PSEXC-CO=-0.086°, dPSECO=0.515°. A Bayesian paired-samples t-test comparing dPSE in both sessions was consistent with physical-driven adaptation: dPSEIL<dPSECO, BF-0=7.530. Therefore, the absence of a global-driven TAE in the Illusion session was not simply because the global pattern of orientation was insufficient to induce TAE – rather, the illusory (but not the physical) global pattern was insufficient to induce TAE.

The overall predominance of global-driven TAE in the Control session, despite presentation of the uniform pattern for only ∼27% of time, may be related to a putatively stronger adaptation during this time due to extra-classical receptive field effects exerted by the global surround on the adapting Gabor when the latter takes the global orientation (iso-orientation surround suppression) [6]. Whatever the contribution of this effect, it acts differently on physical compared to illusory iso-orientation, in the manner expected for low-level processing of the former, but not the latter

### Time-dependent effect

Overall, responses in the Illusion and Control session fit the hypothesis that TAE under UI is driven by physical, and *not* illusory orientation. However, in the Illusion session UI is perceived during only ∼27% of pattern exposure, on average. Thus, it could be argued that a global, illusion-driven TAE might have been present, but undetected in the overall results, -overshadowed by the local-driven TAE at times when UI is not perceived. This possibility seems unlikely, because responses in the Control session (with uniformity also presented ∼27% of time) *do* show an influence of the global-driven TAE. Thus, such a possibility could only hold if the TAE driven by illusory orientation was weaker than that caused by physical orientation. To rule out this possibility, we examined the data from the Illusion sessions for evidence of exposure time-dependency of the TAE magnitude. Since the TAE is time-dependent [7], if illusion-driven adaptation was in fact present, we should find evidence for a shift toward more global/less local TAE with longer times of perceived uniformity.

#### Illusion session

If the TAE is driven *only* by physical orientation, in the Illusion session we should expect independence from time of perceived uniformity. Conversely, if the perceived orientation under UI causes adaptation, the response pattern should shift from predominantly local-driven towards more global-driven for longer time of perceived uniformity. We can assess this potential shift by examining dPSE. As stated above, negative dPSE indicates predominance of local-driven TAE and positive dPSE global-driven TAE. Thus, in the presence of illusion-driven adaptation, dPSE should correlate positively with time of perceived uniformity.

As time measure, we employed the proportion of perceived uniformity (over time of pattern presentation), for conveying the balance between local and (putative) global effects.

For assessing time-dependency within each participant’s data, blocks were classified according to time of perceived uniformity with respect to each participant’s median and dPSE was computed for each category (termed t and T+). Because not every participant had both CX and XC blocks in each category, this led to the loss of eight participants (therefore N=17).

The average proportion of perceived uniformity in t-and T+ was 21.63% and 35.66%, respectively (35.11 and 58.11 seconds). There was a local-driven TAE in both categories: dPSEt-=-0.661o, dPSET+=-1.436o (Figure 3C). According to a Bayesian paired-samples t-test, there was moderate evidence against a positive association with time (dPSEt-<dPSET+): BF-0=0.142.

**Figure 3.**
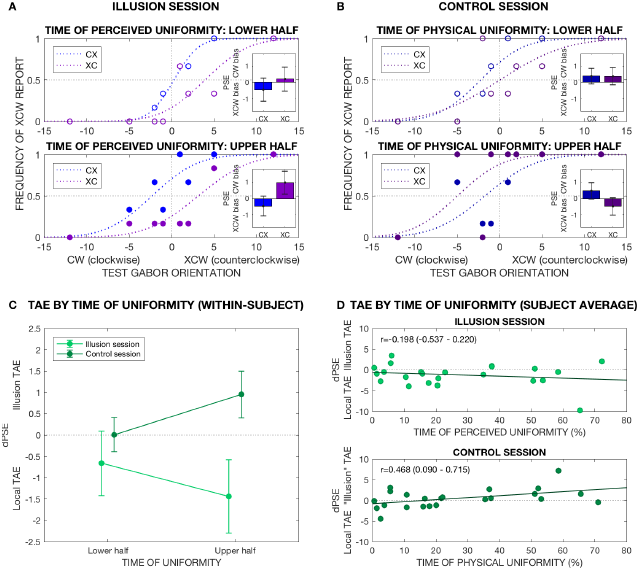
TAE by time of uniformity. **3A – C** classify blocks into lower and upper half according to each participant’s median time of perceived (Illusion session) or physical (Control session) uniformity. **3A-B** depict responses by adapting condition, split by time of uniformity in the Illusion (3A) and Control (3B) session. The main figures present a representative participant’s data; the insets show the sample’s average PSE. In the Illusion session, PSEs indicate local-driven TAE regardless of time of perceived uniformity, while in the Control session the effect shifts to global-driven for longer times of presented physical uniformity. **3C** depicts average dPSE per time of uniformity and session, showing the direction of the TAE in each case. **3D** presents the correlation between individual average proportion of perceived or physical uniformity and dPSE in the Illusion and Control session, respectively. Only in the control session there is a positive correlation with time, as expected from an effect of physical, but not illusory uniformity on the direction of the TAE.

Subsequently we analysed the bivariate correlation between each participant’s average time of perceived uniformity and dPSE (without loss of participants in this case;Figure 3D). Pearson’s correlation coefficient and 95% credible intervals were r=-0.199 (−0.537 – 0.220), with moderate evidence against a positive correlation: BF+0=0.146.

In summary, evidence opposed any positive association between time of perceived uniformity and a trend toward more global-driven TAE, thus opposing predictions expected for illusion-based adaptation.

#### Control session

For the Control session we performed analogous analyses as for the Illusion session, but with time of physical instead of perceived uniformity.

Since global uniformity is a physical stimulus in this session, a time-dependent shift from local to global-driven TAE should be expected regardless of the capacity of illusory orientation to induce a TAE. Thus, this Control session acts as a sanity check to rule out that the failure to find time-dependency in the Illusion session was simply due to insufficient exposure to the global pattern - even in the cases of longest time of uniformity.

For within-subject dependency (Figure 3C) we considered categories t- and T+ based on individual median time of physical uniformity: average time of uniformity per category was 21.42% and 35.74% (31.66 and 52.72 seconds), respectively. As in the illusion session, computing dPSE for both categories led to loss of participants (N=16). Evidence for time-dependency (dPSEt-<dPSET) was only anecdotal: dPSEt-=-0.099o (local TAE), dPSET+=0.989o (global TAE), paired-samples t-test BF-0=1.756.

The correlation between individual average time of physical uniformity and dPSE (Figure 3D) rendered r=0.468 (95% credible intervals 0.090 – 0.715), with moderate evidence for a positive correlation: BF+0=5.540.

In summary, physical uniformity presented for durations equivalent to the reported illusory uniformity was sufficient to observe a shift towards a global-driven TAE.

## Conclusions

The Uniformity Illusion (UI) is a striking phenomenon in which experience across the whole visual field is modified by higher-precision foveal information, yet its underlying mechanisms remain unknown. Using a version of UI with oriented Gabor patches, we found that UI does not produce an orientation adaptation after-effect consistent with the illusory percept. Instead, orientation after-effects only ever followed the (local) physically presented orientation. This suggests that the UI arises from higher-level (higher than primary visual cortex) perceptual processes. It has been suggested that the UI may be the result of predictive processing operations in the visual hierarchy [3]. In a strong predictive-coding interpretation [8], perception of uniformity occurs because high precision central (foveal) information feeds back through the visual hierarchy to alter predictions in lower layers in the processing hierarchy, changing peripheral perception. Our results suggest that if UI does result from such sensory operations, the locus of influence of the feedback does not reach primary visual cortex, as illusory uniformity produced no measurable adaptation effect. It remains an open question whether the absence of a low-level processing effect associated with UI generalizes to other instances of perceptual uniformity (like spatial frequency, density, or colour), and further, precisely what the perceptual mechanisms are that underlie foveal-peripheral integration, as demonstrated by UI, that are central to naturalistic visual experience. However, our results clearly demonstrate that whatever these mechanisms are, they do not appear to alter low-level neural coding in vision.

## ACKNOWLEDGEMENTS

MS-P is supported by a doctoral scholarship from the Dr Mortimer and Theresa Sackler Foundation and the School of Engineering and Informatics, University of Sussex. WR is supported by the EU FET Proactive grant TIMESTORM: Mind and Time: Investigation of the Temporal Traits of Human-Machine Convergence. AKS is grateful for support to the Dr. Mortimer and Theresa Sackler Foundation and to the Canadian Institute for Advanced Research, Azrieli Programme on Mind, Brain and Consciousness.

## REFERENCES

1. Cohen, M.A., D.C. Dennett, and N. Kanwisher, What is the Bandwidth of Perceptual Experience? Trends in Cognitive Sciences, 2016. 20(5): p. 324–335.

2. Haun, A.M., et al., Are we underestimating the richness of visual experience Neuroscience of Consciousness, 2017: p. 1–4.

3. Otten, M., et al., The Uniformity Illusion: Central Stimuli Can Determine Peripheral Perception. Psychological Science, 2016: p. 1–13.

4. Gibson, J.J. and M. Radner, Adaptation, after-effect and contrast in the perception of tilted lines.Journal of Experimental Psychology, 1937. 12: p. 453–467.

5. Knapen, T., et al., The reference frame of the tilt aftereffect. Journal of Vision, 2010. 10(1): p. 8, 1–13.

6. Gheorghiu, E., F.A.A. Kingdom, and N. Petkov, Contextual modulation as de-texturizer. Vision Research, 2014. 104(2014): p. 12–23.

7. Patterson, C.A., S.C. Wissig, and A. Kohn, Distinct Effects of Brief and Prolonged Adaptation on Orientation Tuning in Primary Visual Cortex. The Journal of Neuroscience, 2013. 33(2): p. 532–543.

8. Friston, K.J., Prediction, perception and agency. International Journal of Psychophysiology, 2012. 83(2): p. 248–252.

